# Multiple Wnts act synergistically to induce Chk1/Grapes expression and mediate G2 arrest in Drosophila tracheoblasts

**DOI:** 10.1101/2020.06.01.127266

**Authors:** Amrutha Kizhedathu, Rose Sebastian Kunnappallil, Archit V Bagul, Puja Verma, Arjun Guha

## Abstract

Larval tracheae of Drosophila harbor progenitors of the adult tracheal system (tracheoblasts). Thoracic tracheoblasts are arrested in the G2 phase of the cell cycle in an ATR (mei-41)-Checkpoint Kinase1 (grapes, Chk1) dependent manner prior to mitotic re-entry. Here we investigate developmental regulation of Chk1 activation. We report that Wnt signaling is high in tracheoblasts and is necessary for high levels of activated (phosphorylated) Chk1. We find that canonical Wnt signaling facilitates this by transcriptional upregulation of Chk1 expression in cells that have ATR kinase activity. Wnt signalling is dependent on four Wnts (Wg, Wnt5, 6,10) that are expressed at high levels in arrested tracheoblasts and downregulated at mitotic re-entry. Interestingly, none of the Wnts are dispensable and act synergistically to induce Chk1. Finally, we show that downregulation of Wnt signalling and Chk1 expression leads to mitotic re-entry and the concomitant upregulation of Dpp signalling, driving tracheoblast proliferation.

## INTRODUCTION

The development of a single cell into an organism depends on cells dividing at the appropriate time and place. This requires that cells remain mitotically arrested for a period and resume cycling thereafter. Mitotically competent cells are known to pause either in the G1/G0(Cheung and Rando, 2013) or G2 (Bouldin and Kimelman, 2014) phases of the cell cycle. In this study, we investigate the mechanisms that regulate developmental G2 arrest.

Mechanisms regulating developmental G2 arrest have been studied in Drosophila (Johnston and Edgar, 1998, Ayeni et al., 2016, Otsuki and Brand, 2018) and more recently in Zebrafish (Nguyen et al., 2017). The transition from G2 to M is regulated by an evolutionarily conserved mechanism. Mitotic entry is dependent on the activation of the Cyclin-dependant kinase 1 (Cdk1/Cdc2)-Cyclin B complex, which requires dephosphorylation of Cdc2 by the phosphatase Cdc25. Upon activation, the Cdc2-Cyclin B complex phosphorylates substrates in the cytoplasm and nucleus and prepares the cell for mitosis (Bouldin and Kimelman, 2014). An emerging theme in studies of developmental G2 arrest is that cells paused in G2 lack the essential drivers of G2-M such as Cdc25 and Cyclin B respectively.

Our studies on the mechanism for G2 arrest have focused on the progenitors of the thoracic tracheal (respiratory) system of adult Drosophila (tracheoblasts). Tracheoblasts arrest in G2 for a period of ∼56 h during larval life and then enter mitosis prior to pupariation. The tracheoblasts and the tracheae they comprise grow in size while arrested (increase ∼11 fold in volume) and undergo as rapid size-reductive divisions thereafter (cell division time ∼10 h)(Kizhedathu et al., 2018). Interestingly, tracheoblasts in G2 express all the drivers necessary for G2-M and utilize a distinct mechanism for G2 arrest. We have shown previously that tracheoblasts co-opt the ATR (mei-41)/Chk1(Grapes) DNA damage checkpoint pathway and thereby arrest in G2 (Kizhedathu et al., 2018). Loss of ATR/Chk1 resulted in precocious mitotic re-entry. Moreover, we found that the unusual juxtaposition of positive and negative regulators of G2-M in tracheoblasts was necessary for growth of the cells and the tracheae they comprise. The mechanism by which the ATR/Chk1 axis is controlled in tracheoblasts was not clear from this study. Although the activation of ATR/Chk1 has typically been associated with DNA damage, there was no evidence for DNA damage.

Developmental signals have been shown to regulate G2 arrest by controlling expression of the cell cycle machinery necessary for G2-M. A pulse of Drososphila steroid hormone Ecdysone has been shown to trigger the expression of *Stg* mRNA in histoblasts leading to mitotic re-entry(Ninov et al., 2009). Similarly, Notch and Wnt signalling pathways repress the expression of *Stg* in neural precursors in the wing imaginal disc (Johnston and Edgar, 1998). Insulin signalling in NSCs represses Tribbles expression leading to the upregulation of Stg and resumption of mitotic cycling (Otsuki and Brand, 2018). Here we investigate the developmental signals regulate the activation of the ATR/Chk1 axis in tracheoblasts.

To identify candidate signals, we knocked down the components of nine developmental signalling pathways that regulate cell proliferation and examined whether these perturbations resulted in precocious mitotic re-entry. We found that perturbations in Wnt signalling led to premature cell division akin to Chk1 mutants and in turn examined the roles of the Wnt signalling pathway in the regulation of the ATR/Chk1 pathway.

## RESULTS

### Wnt Signalling is required to maintain G2 arrest in Tr2 tracheoblasts

The tracheal branches of the second thoracic metamere (Tr2) in the Drosophila larva are comprised of cells that also serve as the progenitors of the thoracic tracheal system of the adult animal. These tracheoblasts remain quiescent through most of larval life and rekindle a mitotic program prior to pupariation. Tracheoblasts in the Dorsal Trunk (DT) in Tr2 enter larval life in G1, transition from G1 to S to G2 in the first larval instar (L1) and remain arrested in G2 from second instar (L2) till 32-40 h into third larval instar (L3) whereupon they rekindle a mitotic program (Figure 1A) (Kizhedathu et al., 2018). These cells express high levels of Chk1 mRNA and phosphorylated Chk1 (pChk1) during the period in which they are paused in G2 and downregulate levels of Chk1 mRNA and pChk1 upon mitotic re-entry. Loss of Chk1 leads to precocious proliferation by 16-24 h L3, ∼24 h earlier than expected (Figure 1B).

**Figure 1:**
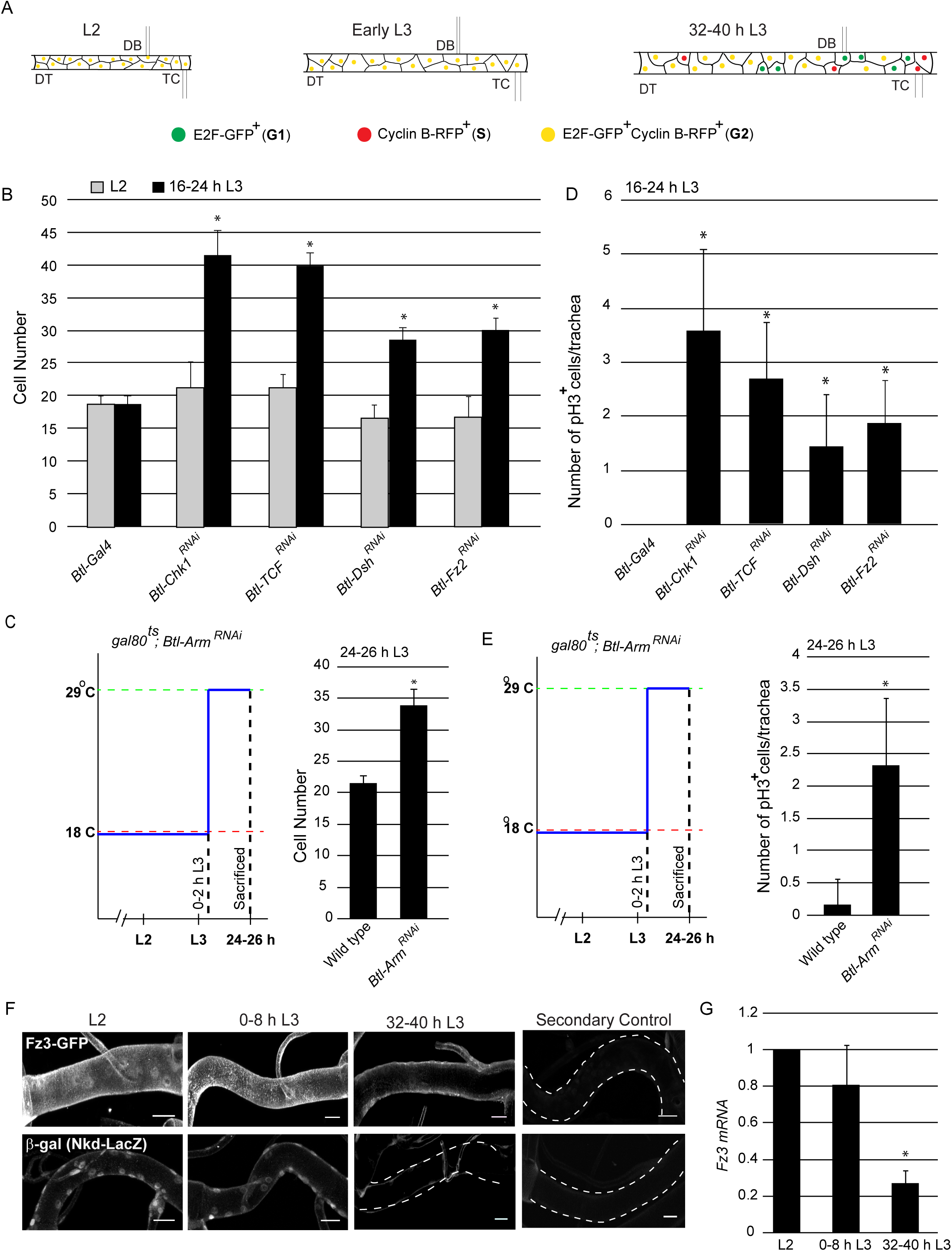
Wnt signalling is required for G2 arrest in Tr2 tracheoblasts. **(A)** Cartoon representing the cell cycle phasing of cells in Tr2 DT at different larval stages based on FUCCI. **(B-C)** Impact of knockdown of *Chk1* and components of the Wnt signalling pathway on cell numbers in Tr2 DT. **(B)** Graph shows cells numbers in wild type (*Btl-Gal4*), *Btl-Chk1*^*RNAi*^, and *Btl-*Wnt pathway components ^RNAi^ at L2 and 16-24 h L3. **(C)** Graph shows cell numbers in wild type (*tub-Gal80*^*ts*^*/+; UAS-Arm*^*RNAi*^/*Tb)* and Armadillo mutants (*tub-Gal80*^*ts*^*/+; Btl-Gal4/UAS-Arm*^*RNAi*^) at 24-26h L3. Larvae were grown at 18°C and transferred to 29°C at 0-2 h into L3. **(D-E)** Impact of reduction in levels of *Chk1* or in levels of different components of the Wnt signalling pathway on mitotic indices in Tr2 DT. **(D)** Graph shows frequency of pH3^+^ nuclei at 16-24 h L3 in wild type (*Btl-Gal4*), *Chk1* mutant, and Wnt pathway mutants **(E)** Graph shows the frequency of pH3^+^ nuclei in wild type (*tub-Gal80*^*ts*^*/+; UAS-Arm*^*RNAi*^/*Tb)* and Armadillo mutants (*tub-Gal80*^*ts*^*/+; Btl-Gal4/UAS-Arm*^*RNAi*^) at 24-26h L3. Larvae were grown as stated above. **(F)** Expression pattern of Wnt reporters *Fz3*-GFP and *Nkd*-LacZ in Tr2 DT at different larval stages. Panels show immunostaining for GFP in *Fz3*-GFP (top pane) and β-Gal in *Nkd*-LacZ (Bottom panel) and their respective secondary controls **(G)** Quantitative PCR analysis of *Fz3* mRNA levels in micro-dissected Tr2 DT fragments at different stages. Graph shows fold change in mRNA levels with respect to L2 (n=3 experiments, n>15 Tr2 DT fragments/condition/experiment, mean±standard deviation). DT=Dorsal Trunk, DB=Dorsal Branch, TC=Transverse Connective. Scale bar = 20 µm (mean±standard deviation, n>7 tracheae) Student’s paired t-test: *p<0.05.

To explore the possibility that developmental signals control the activation of the ATR/Chk1 axis in tracheoblasts, we knocked-down expression of essential transducers of the EGF, FGF, insulin/PI3K, Hedgehog, JAK/STAT, Ecdysone, Notch, Wnt and Dpp signalling pathways in the trachea by RNA interference (under control of *Breathless (Btl)*-*Gal4*) and determined if these perturbations led to precocious mitotic re-entry at 16-24 h in L3. Of the abovementioned signalling pathways, reduced expression of components of Wnt (Figure 1) pathways resulted in precocious proliferation at 16-24 h L3. These findings led us to examine more carefully the roles of Wnt in the activation ATR/Chk1 axis in tracheoblasts.

The Wingless/Wnt signalling pathway regulates many different developmental processes in metazoans. Wnt genes encode secreted proteins that can act as signalling molecules and morphogens. The Wnt pathway is activated upon the binding of Wnt ligands to Frizzled (Fz) receptors. Ligand-receptor engagement leads to phosphorylation of Dishevelled (Dsh) that in turn antagonizes an intracellular complex that targets β-Catenin for degradation. The stabilization of β-Catenin facilitates its translocation to the nucleus where it binds the TCF/LEF family of transcription factors and initiates the transcription of downstream targets (Bejsovec, 2006). Apart from this “canonical” mechanism there are “non-canonical” mechanisms for Wnt signalling that do not require β-Catenin stabilization (Zhan et al., 2017, Swarup and Verheyen, 2012).

To characterize the role of Wnt signalling in G2 arrest in tracheoblasts, we knocked-down the abovementioned components of the pathway in the trachea by RNA interference and examined how these impacted timing of mitotic re-entry. To determine whether perturbations in Wnt signalling led to precocious mitotic entry we used two assays. First, we quantified the numbers of tracheoblasts in Tr2 DT at L2 and at 16-24 h L3. Second, we quantified the frequencies of phospho-histone H3^+^ (pH3^+^) mitotic figures in Tr2 DT at 16-24 h L3. Knockdown of *Fz2, Dsh and TCF (Pangolin)* did not affect numbers of tracheoblasts in L2 but led to increased numbers at 16-24 h L3 (Figure 1B, n > 7 tracheae per timepoint here and in all subsequent figures showing cell frequencies). In our hands, the knockdown of Drosophila β-Catenin (*Armadillo (Arm)*) in the tracheal system from embryonic stages resulted in embryonic lethality. Thus, to characterize the role of β-Catenin in tracheoblasts at larval stages we knocked-down *Arm* expression in the trachea specifically at larval stages using the temperature sensitive Gal4-UAS-gal-80^ts^ system (see schematic in Figure 1C). Briefly, control (*tub-Gal80*^*ts*^*/+; UAS-Arm*^*RNAi*^/*Tb)* and *Btl*-*Arm*^*RNAi*^ (*tub-Gal80*^*ts*^*/+; Btl-Gal4/UAS-Arm*^*RNAi*^) animals were grown at 18°C till 0-2 h L3 and moved to 29°C for 24 h. The animals were then sacrificed at 24-26 h L3 and the cell numbers in Tr2 DT were counted. Knockdown of *Arm* did not perturb numbers of cells in Tr2 DT in L2 but resulted in increased numbers of cells at 24 h L3 (Figure 1C, n > 7 tracheae per condition). Apart from quantifying cell numbers at L2 and early L3, we assayed mitotic activity by immunostaining with an antibody against phosphorylated Histone H3 (pH3), a marker for cells undergoing mitosis. We found increased frequencies of pH3^+^ figures at early L3 in all backgrounds in which we observed increased numbers of cells at early L3 (Figure 1D-E). We also compared the timecourse of cell proliferation in L3 in wild type, Chk1 and Wnt pathway mutants. We found that in both Chk1 and Wnt mutant cells enter mitosis precociously and continue cell division thereafter (Figure 1 Figure Supplement 1). Together, these data show that loss of Wnt signalling leads to precocious mitotic re-entry and proliferation.

In light of the findings that Wnt signalling is required for G2 arrest, we next investigated the spatiotemporal pattern of Wnt signalling in Tr2 DT during larval stages. For this we examined the expression of three reporters for Wnt signalling (*Frizzled3 (Fz3)*-GFP and *Naked* (Nkd)-LacZ) at L2, 0-8 h L3 and 32-40 h L3 (Sivasankaran et al., 2000, Xu et al., 2018, Tian et al., 2016). Immunostaining using anti-GFP and anti-β-gal antisera revealed that levels of expression of *Fz3*-GFP and *Nkd*-LacZ were high at L2 and 0-8 h L3 and significantly lower at 32-40 h into L3 (Figure 1F, n = 6 tracheae per condition per experiment, n=2). We also measured the expression of *Fz3* mRNA in Tr2 DT by quantitative real time PCR (qPCR). qPCR analysis showed that the expression of *Fz3* mRNA is high in L2 and early L3 and significantly lower at 32-40 h L3 (Figure 1G, n=3). These data show that canonical Wnt signalling is active in cells while they are paused in G2 and downregulated upon mitotic re-entry.

### pChk1 levels are reduced in Wnt pathway mutants

Since perturbations in Wnt signalling led to precocious proliferation akin to loss of Chk1, next we examined the role of Wnt pathway with regard to Chk1 activation (phosphorylated Chk1, pChk1). Immunostaining Tr2 DT in wild type animals with an antibody against pChk1 has shown that levels are high in tracheoblasts in L2 and early L3 and significantly lower at 32-40 h L3 (Kizhedathu et al., 2018). pChk1 immunostaining in *Btl*-*TCF*^*RNAi*^ at L2 showed that the levels of pChk1 were dramatically reduced when compared to wild type at this stage and comparable to background (Figure 2A, n = 6–8 tracheae per condition per experiment, n = 3). We inferred that loss of Wnt signalling leads to a reduction in pChk1 levels. These data showed that Wnt pathway indeed regulates Chk1 activation.

**Figure 2:**
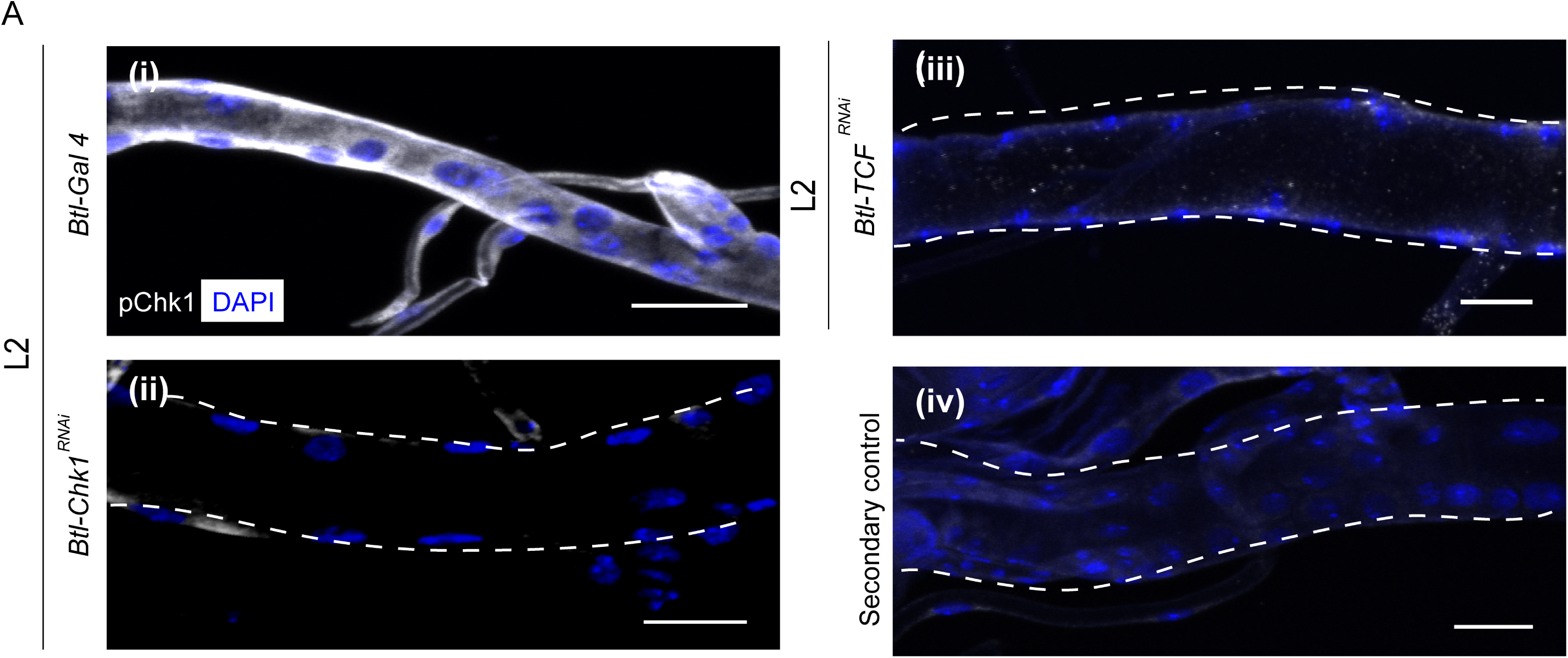
Phosphorylated Chk1 levels are diminished in the absence of Wnt signalling. **(A)** Activated Chk1 (phospho-Chk1^Ser345^, pChk1) immunostaining (white) in Tr2 DT in Chk1 mutants and Wnt pathway mutants. pChk1 immunostaining in Tr2 DT in **(i)** wild type (*Btl-Gal4*), **(ii)** *Btl-Chk1*^*RNAi*^ **(iii)** *Btl-TCF*^*RNAi*^ at L2 and **(iv)** wild type treated with secondary antibody alone. Scale bar = 20 µm

### Wnt signalling upregulates Chk1 expression

Our earlier studies have shown that levels of pChk1 and Chk1 mRNA are correlated in Tr2 DT. In other words, Chk1 mRNA levels are also high in L2 and early L3 and diminish at 32-40 h L3. This led us to hypothesize that Wnt signalling facilitates high levels of pChk1 via induction of high levels of Chk1 expression. To investigate the relationship between Wnt signalling and Chk1 expression, we examined Chk1 promoter activity in wild type and Wnt pathway mutants using an enhancer trap line *(Chk1-LacZ).* Anti-β-gal immunostaining of tracheae from *Chk1-LacZ* animals at different larval stages showed that β-gal expression is high in L2 and early L3 and declines at 32-40 h into L3 (Figure 3A, n = 6 tracheae per condition per experiment, n = 2). We then analysed expression of the reporter in *Btl-TCF*^*RNAi*^ animals at L2. We found that β-gal expression was considerably diminished in L2 (Figure 3A, n = 6 tracheae per condition per experiment, n = 2). We also quantified the levels of Chk1 mRNA in microdissected Tr2 fragments from wild type, *Btl-TCF*^*RNAi*^ and *Btl-ArmS10 (*an activated form of *Armadillo* (*Btl-ArmS10*)) at different larval stages using quantitative real-time PCR (qPCR). The loss of Wnt signalling (*Btl-TCF*^*RNAi*^) led to a reduction in *Chk1* levels at L2 and early L3; the levels of expression at these stages were comparable to 32-40 h L3. Conversely, overexpression of *ArmS10* led to an increase in the level of Chk1 transcript (Figure 3B, ≥15 tracheal fragments per timepoint per experiment, n = 3 experiments). Together, the analysis of Chk1 mRNA levels and *Chk1*-*LacZ* show that Wnt signalling is required for high levels of *Chk1* expression in arrested cells in Tr2 DT.

**Figure 3:**
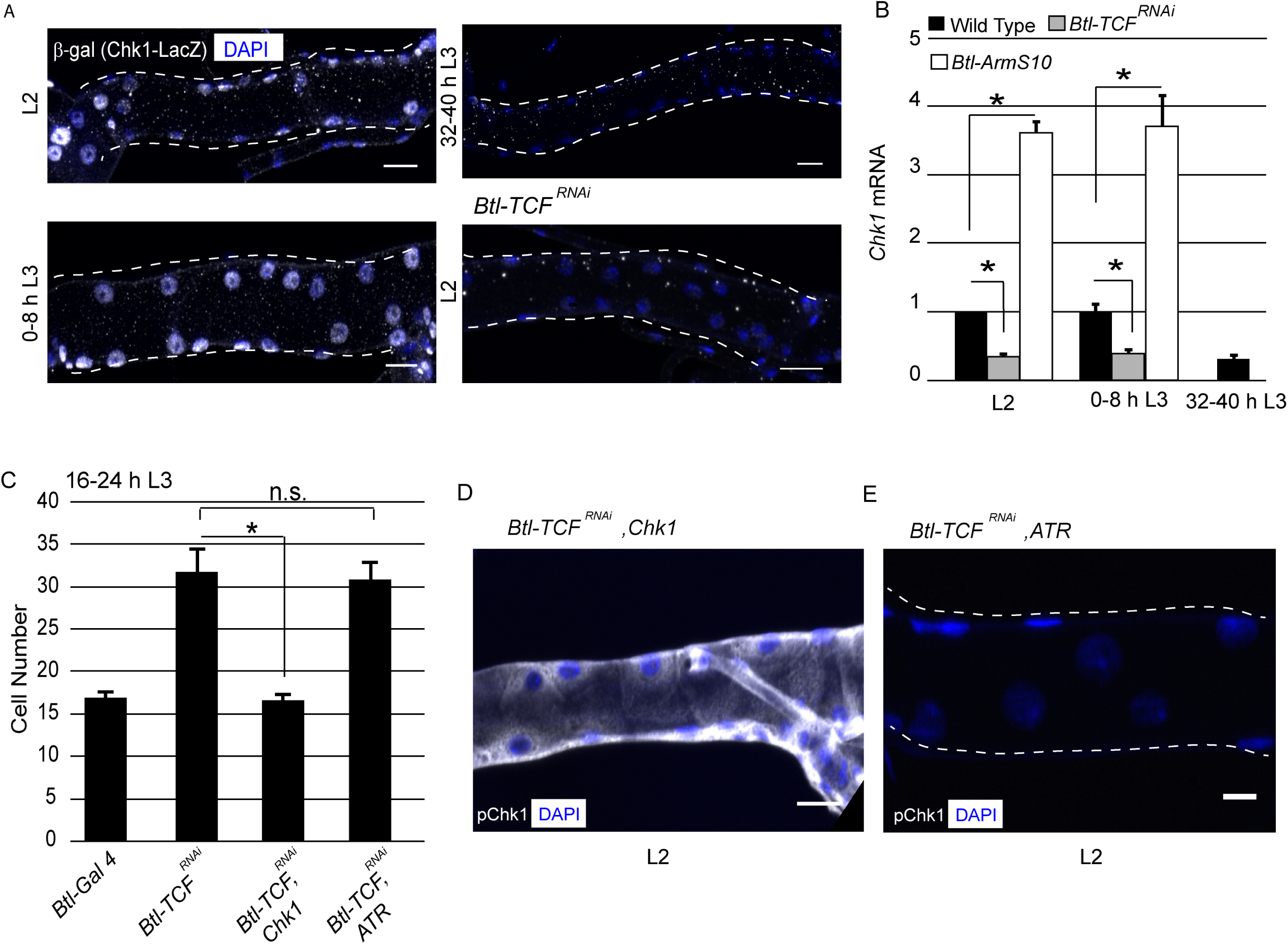
Wnt signalling regulates Chk1 transcription. **(A)** Analysis of *Chk1-*LacZ expression in Tr2 DT in wild type and Wnt pathway mutants. β-Gal immunostaining in larvae expressing *Chk1-*LacZ at L2, 0-8 h L3 and 32-40 h L3 and in *Btl-GAL4/ Chk1-LacZ; TCF*^*RNAi*^*/+* at L2. **(B)** Quantitative PCR analysis of *Chk1* mRNA levels in micro-dissected Tr2 DT fragments at different stages. Graph shows fold change in *Chk1* mRNA in Tr2 DT fragments from wild type (*Btl-Gal4*), Wnt pathway loss-of-function (*Btl-Gal4/+; UAS-TCF*^*RNAi*^*/+)* and Wnt pathway gain-of-function (*UAS-ArmS10/+; Btl-Gal4/+)* larvae. Fold change has been represented with respect to L2 (n=3 experiments, n>15 Tr2 DT fragments/condition/experiment, mean±standard deviation). **(C)** Impact of overexpression of *Chk1* and *ATR* in TCF mutants. Graph shows cell numbers in wild type (*Btl-Gal4, UAS-FUCCI/Cyo GFP; +/Tb*), *Btl-TCF*^*RNAi*^ *(Btl-Gal4, UAS-FUCCI/+; UAS-TCF*^*RNAi*^*/+*), *Btl-TCF*^*RNAi*^, *Chk1* (*Btl-Gal4, UAS-FUCCI/+; UAS-TCF*^*RNAi*^*/ UAS-Chk1) and Btl-TCF*^*RNAi*^, *ATR* (*Btl-Gal4, UAS-FUCCI/+; UAS-TCF*^*RNAi*^*/ UAS-ATR)* at 16-24 h L3. **(D)** Impact of reduction of *TCF* and overexpression of *Chk1* on levels of pChk1 in Tr2 DT. pChk1 immunostaining (white) in *Btl-TCF*^*RNAi*^, *UAS-Chk1* (*Btl-Gal4, UAS-FUCCI/+; UAS-TCF*^*RNAi*^*/ UAS-Chk1)* animals at L2 (This image has been acquired at a lower laser power and gain setting compared to Figure 2A to prevent saturation of white pixels). **(E)** Impact of reduction of *TCF* and overexpression of *ATR* on levels of pChk1 in Tr2 DT. pChk1 immunostaining (white) in *Btl-TCF*^*RNAi*^, *UAS-ATR* (*Btl-Gal4, UAS-FUCCI/+; UAS-TCF*^*RNAi*^*/ UAS-ATR)* animals at L2 (mean values±standard deviation, n>7 tracheae) Scale bar = 20 µm. Student’s paired t-test: *p<0.05

To test the hypothesis that Wnt signalling regulates G2 arrest via control of Chk1 expression, we overexpressed *Chk1* or *ATR* in *Btl-TCF*^*RNAi*^ animals and determined if this prevented precocious proliferation and restored levels of pChk1. Examination of cell numbers in Tr2 DT at 16-24 h L3 in Wild type, *Btl-TCF*^*RNAi*^, *Btl-TCF*^*RNAi*^, *Chk1* and *Btl-TCF*^*RNAi*^, *ATR* expressing animals showed that Chk1 overexpression did indeed prevent premature mitotic re-entry (Figure 3C). The overexpression of ATR however did not prevent precocious proliferation (Figure 3C). Immunostaining against pChk1 revealed that the levels of active Chk1 in *Btl-TCF*^*RNAi*^, *Chk1* animals was comparable to wild type. In contrast, the levels of pChk1 were not restored in *Btl-TCF*^*RNAi*^, *ATR* expressing animals and were comparable to *Btl-Chk1*^*RNAi*^ animals at the same stage (Figure 3D-E, n = 6 tracheae per experiment, n = 2). These data support the hypothesis that Wnt dependent regulation of Chk1 expression is necessary for G2 arrest in Tr2 DT.

### *Wg, Wnt5, Wnt6* and *Wnt10* are required for Chk1 expression

Since Wnt signalling is high when the cells are arrested and diminish when the cells start dividing, we hypothesized that the ligands that activate Wnt signalling would also follow the same pattern. To test this hypothesis, we measured the RNA levels of the various Wnt ligands in microdissected Tr2 fragments using qPCR. The analysis revealed that mRNA levels of 4 Wnt ligands – *Wg, Wnt5, Wnt6* and *Wnt10* - are high in L2 and 0-8 h L3 and low at 32-40 h L3 (Figure 4A, ≥15 tracheal fragments per timepoint per experiment, n = 3 experiments). *Wnt2, Wnt4* and *WntD* were undetectable in the tracheae (data not shown). To further validate this analysis, we performed single molecule FISH (smFISH) on the trachea for *Wg, Wnt5, Wnt6* and *Wnt10*. The signal consisted of prominent nuclear spots representing the transcription sites and faint spots in the cytoplasm marking accumulated mRNA. We observed that at L2 (Figure 4B) and 0-8 h L3 (data not shown), we could detect prominent nuclear spots (Figure 4B Arrowheads) and faint cytoplasmic signal. We did not detect either at 32-40 h L3. Together, this data suggests that *Wg, Wnt5, Wnt6* and *Wnt10* are expressed in the trachea when the cells are arrested in G2 and diminish when the cells enter mitosis.

**Figure 4:**
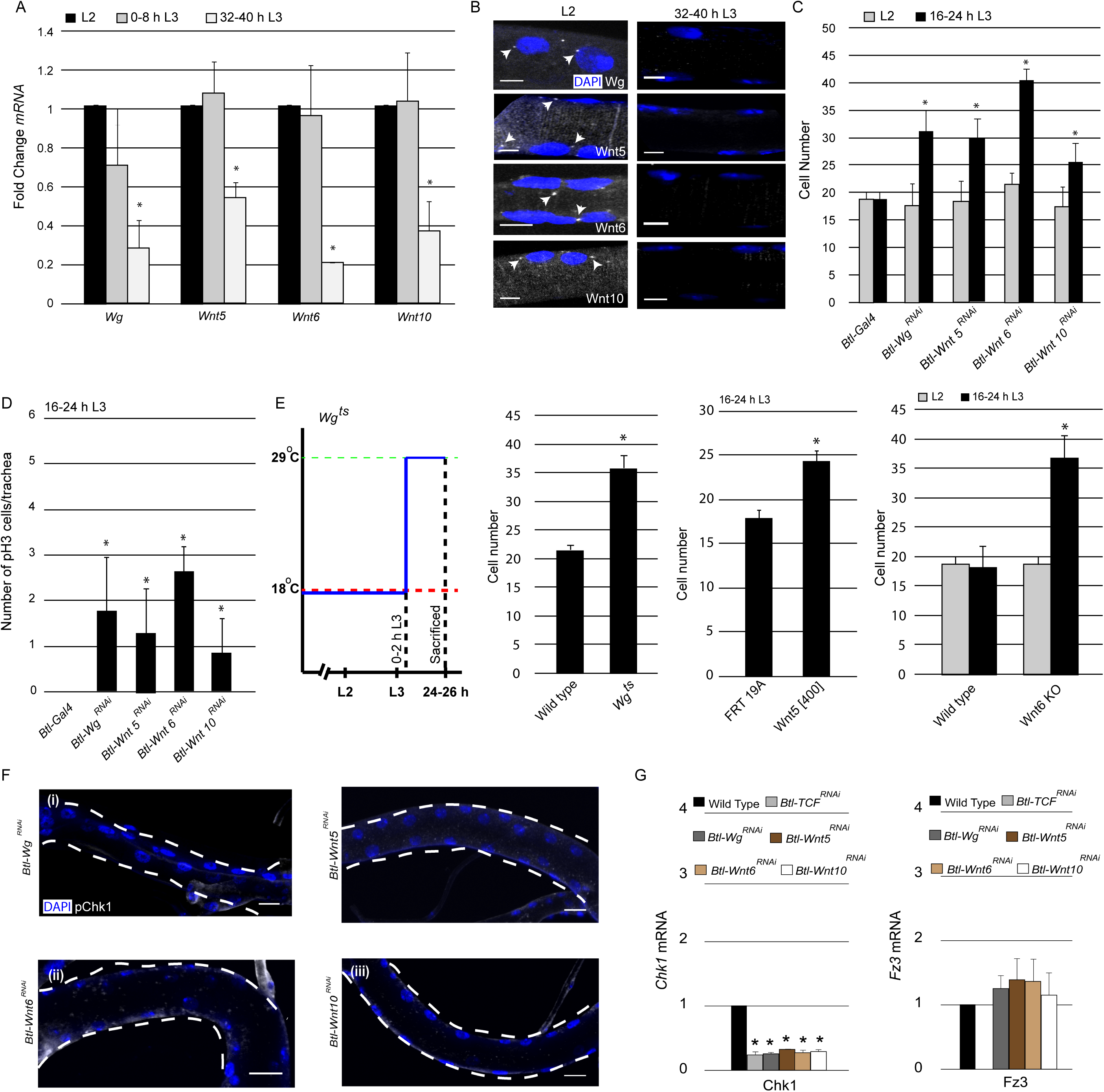
Wg, Wnt5, Wnt6 and Wnt10 are required to maintain Chk1 expression. **(A)** Quantitative PCR analysis for levels of *Wg, Wnt5, Wnt6 and Wnt10* mRNA in micro-dissected Tr2 DT fragments at different stages. Graph shows fold change in *Wg, Wnt5, Wnt6 and Wnt10* mRNA in Tr2 DT fragments from wild type (*Btl-Gal4)* larvae at L2, 0-8 h L3 and 32-40 h L3. Fold change has been represented with respect to L2 (n=3 experiments, n>15 Tr2 DT fragments/condition/experiment, mean±standard deviation). **(B)** smFISH for *Wg, Wnt5, Wnt6 and Wnt10* mRNA in Tr2 DT at L2 and 32-40 h L3. Arrowheads point to the sites of mRNA accumulation. (Scale bar = 5 µm) **(C)** Impact of knockdown of components of the Wnt signalling pathway on cell numbers in Tr2 DT. Graph shows cells numbers in wild type (*Btl-Gal4*, Same as Figure 1 B) and *Btl-*Wnt pathway components^RNAi^ at L2 and 16-24 h L3. **(D)** Impact of reduction in levels of different components of the Wnt signalling pathway on mitotic indices in Tr2 DT. Graph shows frequency of pH3^+^ nuclei at 16-24 h L3 in wild type (*Btl-Gal4*, Same as Figure 1 D) and Wnt pathway mutants **(E)** Impact of expression of mutant *Wg, Wnt5 and Wnt6* on cell numbers in Tr2 DT. Graphs show cell numbers in wild type and *Wg* mutants ((*Wg*^*ts*^) at 24-26h L3, Larvae were grown at 18°C and transferred to 29°C at 0-2 h into L3) *Wnt5* mutant *(Wnt5[400] FRT19A)* and *Wnt6 (Wnt6 KO)* on cell numbers in Tr2 DT. **(F)** Activated Chk1 (phospho-Chk1^Ser345^, pChk1) immunostaining (white) in Tr2 DT in Wnt pathway mutants. pChk1 immunostaining in Tr2 DT in (i) *Btl-Wg*^*RNAi*^, (ii) *Btl-Wnt5*^*RNAi*^ (iii) *Btl-Wnt6*^*RNAi*^ and *Btl-Wnt10*^*RNAi*^ at L2. **(G)** Quantitative PCR analysis of *Chk1 and Fz3* mRNA levels in micro-dissected Tr2 DT fragments at L2. Graphs show fold change in *Chk1 and Fz3* mRNA in Wild type, *Btl-TCF*^*RNA*^, *Btl-Wg*^*RNAi*^, *Btl-Wnt5*^*RNAi*^, *Btl-Wnt6*^*RNAi*^ and *Btl-Wnt10*^*RNA*^ in Tr2 DT fragments. Fold change has been represented with respect to Wild type (n=3 experiments, n>15 Tr2 DT fragments/condition/experiment, mean±standard deviation). (mean values±standard deviation, n>7 tracheae) Scale bar = 20 µm. Student’s paired t-test: *p<0.05

Next we examined the effect of reduction of the abovementioned components on the timing of mitotic entry in Tr2 DT. For this, we quantified the number of cells in Tr2 DT at L2 and 16-24 h L3 and the frequency of pH3^+^ nuclei in Tr2 DT at 16-24 h L3. Knockdown of *Wg, Wnt5, Wnt6* and *Wnt10* in the trachea led to an increase in cell number at 16-24 h into L3 (Figure 4C). There was also an increase in pH3^+^ figures in these backgrounds at 16-24 h L3 (Figure 4D). We also quantified the cell numbers in *Wg (Wg*^*ts*^)(Swarup and Verheyen, 2012), *Wnt5 (Wnt5[400]*(Fradkin et al., 2004)) and *Wnt6(Wnt6 KO)*(Doumpas et al., 2013) mutants at early L3 to further validate these observations. *Wg*^*ts*^ animals were grown at 18°C till 0-2 h L3 and moved to 29°C for 24 h. The animals were then sacrificed at 24-26 h L3 and the cell numbers in Tr2 DT were counted. We found that though there is no perturbation in cell numbers at L2, there in an increase in cell numbers at 24-26 h L3 (Figure 4E). Quantification of cell numbers at L2 and16-24 h L3 in *Wnt5[400]* and *Wnt6 KO* revealed that there are increased cell numbers in these backgrounds at 16-24 h L3. Taken together, this data suggests *Wg, Wnt5, Wnt6* and *Wnt10* are required to maintain G2 arrest in Tr2 DT.

As knockdown of any one of the ligands led to precocious proliferation, we examined how each of these perturbations impacts levels of pChk1 and *Chk1* mRNA. For this, we examined the levels of pChk1 in Tr2 and also the levels of *Chk1* mRNA by qPCR. Analysis of pChk1 levels in *Btl-Wg*^*RNAi*^, *Btl*-*Wnt5*^*RNA*i^, *Btl*-*Wnt6*^*RNAi*^ and *Btl*-*Wnt10*^*RNAi*^ revealed that the levels of pChk1 in these animals were comparable to levels of pChk1 in *Btl-Chk1*^*RNAi*^ expressing animals (Figure 4F, compare with Figure 2A (ii)). Quantification of *Chk1* mRNA levels in *Btl*-*TCF*^*RNAi*^, *Btl-Wg*^*RNAi*^, *Btl*-*Wnt5*^*RNA*i^, *Btl*-*Wnt6*^*RNAi*^ and *Btl*-*Wnt10*^*RNAi*^ expressing animals showed that the loss of any one of the ligands diminished the levels of mRNA to levels comparable to that of *Btl*-*TCF*^*RNAi*^ (Figure 4G). We also examined the mRNA levels of *Fz3*, a conventional target of Wnt signalling, in animals expressing *Btl-TCF*^*RNAi*^, *Btl-Wg*^*RNAi*^, *Btl*-*Wnt5*^*RNA*i^, *Btl*-*Wnt6*^*RNAi*^ and *Btl*-*Wnt10*^*RNAi*^. While the loss of TCF reduced the levels of Fz3 mRNA to levels below detection, there was no significant change in the levels of Fz3 mRNA levels in the other conditions (Figure 4G). This suggests that all four ligands are required to maintain levels of Chk1 to enforce G2 arrest in Tr2 tracheoblasts.

### Exit from G2 arrest is required for the activation of Dpp signalling

The timecourse of proliferation in Chk1 and Wnt pathway mutants showed that tracheoblast enter mitosis precociously at 16-24 h L3 and continue to proliferate thereafter (Figure1 - Figure Supplement 1). A previous study has identified Dpp (the drosophila homolog of TGF-β) as the signal that promotes proliferation post mitotic re-rentry. Dpp signalling is initiated by the binding of the ligand (Dpp) to the receptor Serine/Threonine Kinase Thickveins (Tkv). This in turn which leads to the phosphorylation of the transcriptional activator Mad. Phosphorylated Mad (pMad) translocates to the nucleus and regulates gene expression. Pertinently, the earlier study on tracheoblasts showed that Dpp signalling (pMad immunostaining) was undetected in cells in G2 arrest and upregulated mid L3 when the cells start dividing (Djabrayan and Casanova, 2016). In light of the proliferation dynamics in Chk1 and Wnt pathway mutants, we examined the levels of Dpp signalling in these mutant backgrounds prior to and post mitotic entry.

Consistent with the previous study, pMAD immunostaining was undetectable in L2 and early L3 (0-8 h L3 and 16-24 h L3) and nuclear localized at 32-40 h L3, once the cells entered mitosis (Figure 5A). Next we assayed for pMad in tracheae from *Btl-Chk1*^*RNAi*^ and *Btl-Wnt6*^*RNAi*^ at both L2 and 16-24 h L3. We did not detect any pMad immunostaining at L2 in either but found abundant nuclear localized signal at 16-24 h L3 in both *Btl-Chk1*^*RNAi*^ and *Btl-Wnt6*^*RNAi*^ (Figure 5B). This suggested that release from G2 arrest leads the concomitant upregulation of Dpp signalling. We then investigated whether the expression of an activated form of *Tkv (Btl-Tkv*^*QD*^) (Djabrayan and Casanova, 2016) in the trachea would result in the ectopic activation of pMad at L3. pMad immunostaining was undetectable in *Btl-Tkv*^*QD*^ animals at L2 (Figure 5C). Taken together, these observations suggest that arrest in G2 prevents the activation of Dpp signalling and that tracheoblasts activate Dpp signalling following exit from G2.

**Figure 5:**
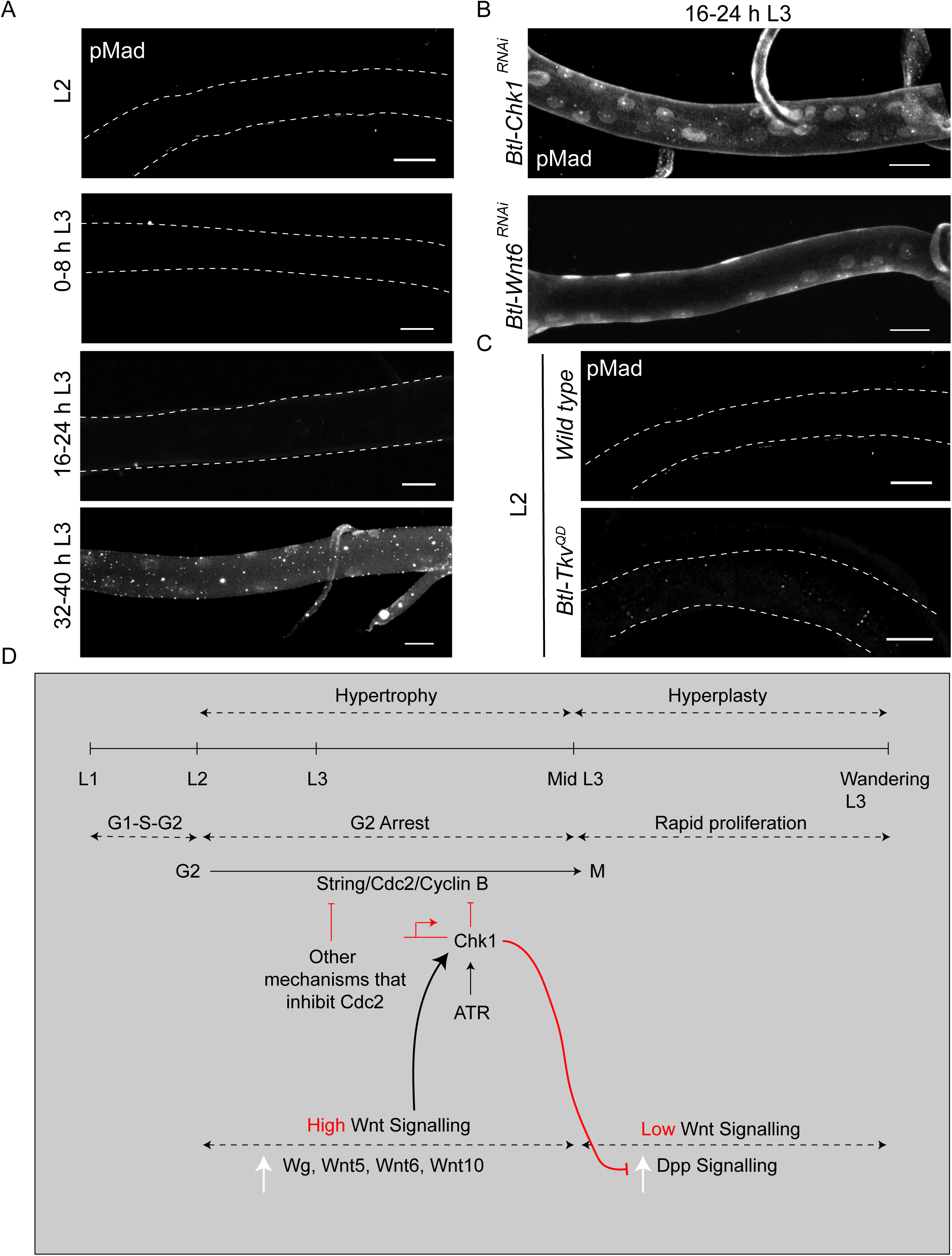
Exit from G2 arrest is required for activation of Dpp signaling. **(A)** phospho-Mad (pMad) immunostaining in wild type Tr2 DT at L2, 0-8 h L3, 16-24 h L3 and 32-40 h L3. **(B)** Impact of reduction of *Chk1* or Wnt pathway components on pMad expression in Tr2 DT. Immunostaining against pMad in Tr2 DT from *Btl-Chk1*^*RNAi*^ and *Btl-Wnt6*^*RNAi*^ animals at 16-24 h L3. **(C)** Impact of overexpression of activated form of Tkv (*Btl-Tkv*^*QD*^) on pMad expression in Tr2 DT. Immunostaining against pMad in Tr2 DT from wild type and *Btl-Tkv*^*QD*^ animals at L2. **(D)** Model for the regulation of G2 arrest by Wnt signalling. Wnt signalling negatively regulates G2-M by transcriptionally upregulating Chk1.Arrest in G2 negatively regulates Dpp signalling, preventing precocious proliferation and allowing for hypertrophic growth of Tr2 DT. Scale bar = 20 µm.

## DISCUSSION

Progenitors of the thoracic tracheal system of adult Drosophila arrest in G2 in a Chk1-dependent manner during larval stages and proliferate thereafter. The goal of this study was to decipher how the activation of Chk1 is developmentally regulated in these cells. Taken together, our findings suggest that Wnt signaling drives high levels of *Chk1* expression in tracheoblasts and that high levels of Chk1 in cells that have a basal level of ATR activity is sufficient to induce G2 arrest. Consistent with the proposed model, studies in yeast (Ford et al., 1994), Drosophila (Bayer et al., 2018) and Xenopus (Kumagai et al., 1998) have shown that the overexpression of Chk1 is sufficient to arrest cells in G2. Our studies also indicate that the developmental regulation of Wnt signalling itself is under transcriptional control. We find that of expression of four Wnt ligands that activate the Wnt pathway, Wg, Wnt5, Wnt6, Wnt 10 are expressed at high levels in tracheoblasts that are in G2 arrest and downregulated at the time the cells enter division.

A fascinating aspect of Wnt signalling in the trachea is that as many as 4 ligands appear to be required; *Wg, Wnt5, Wnt6* and *Wnt10* are all necessary in these cells to maintain G2 arrest. The loss of any one of the ligands leads to a reduction in Chk1 mRNA levels in tracheoblasts comparable with the knockdown of TCF. Interestingly, we also measured how the loss of any of these Wnts or TCF impacted the levels of *Fz3*, a common transcriptional target of canonical Wnt signalling. Our data show that the Wnts act redundantly to regulate Fz3 in the same cells. We hypothesize that the 4 ligands act synergistically to maintain a threshold of Wnt signalling necessary for *Chk1* expression. Consistent with this, we find that overexpression of Wnt5 in Wnt6^RNAi^-expressing animals was able to prevent precocious proliferation at 16-24 h L3 (*Btl*-*Wnt6*^*RNAi*^ at 16-24 h L3 Avg: 32.4 ± 1.2 cells, n=8 tracheae; *Btl*-*Wnt6*^*RNAi*^, *Wnt5* at 16-24 h L3 Avg: 19.1 ± 0.8 cells, n=8 tracheae).

It is noteworthy that *Wnt5*, which in *Drosophila* has been implicated in non-canonical Wnt signalling (Shimizu et al., 2011), facilitates canonical Wnt signalling in tracheoblasts. There is evidence in other systems that non-canonical Wnt ligands can activate canonical Wnt signalling (Fu et al., 2016, Lyons et al., 2004). Preliminary expression profiling of the larval tracheal system has indicated that both Derailed and Doughnut, RYK family receptors that mediate non-canonical Wnt signalling, are expressed (AG, AK unpublished). This suggests that Wnt5 could act either directly or indirectly to facilitate canonical Wnt signalling in tracheoblasts.

Thoracic tracheoblasts exhibit an unusual mode of G2 arrest in that the cells express both drivers of G2-M (Cdc2/Cdk1, CyclinB, Stg) and simultaneously activate the negative regulators of G2-M like Chk1(Kizhedathu et al., 2018). Based on our previous analysis of Chk1 mutant tracheae and subsequent studies we think that the juxtaposition of positive and negative regulators is required for the growth of the tracheoblasts and the tracheae they comprise (see model in Figure 5 and Figure 5, Figure Supplement 1). In light of the role of Wnt signalling in the regulation of Chk1, we have also and examined how Wnt mutants impact growth of the tracheae. Quantification of percentage growth between L2 and 32-40 h L3 revealed that compared to a 268 ± 32% (n = 6 tracheae) growth in length and 247 ± 19% (n = 12 tracheae) growth in width in control animals (Kizhedathu et al., 2018), TCF mutants grew only by 178 ± 8% (n = 6 tracheae) in length and 176 ± 13% (n = 6 tracheae) in width. Thus, consistent with expectation, Wnt signalling is required for optimal growth of the thoracic tracheae during larval stages.

Upon release from G2 arrest, tracheoblasts undergo rapid size-reductive divisions. Cell proliferation is thought to be under the control of Dpp signalling. In light of the finding that precocious mitotic entry leads to sustained proliferation, we investigated the relationship between exit from G2 arrest and the onset of Dpp signalling. The data suggests that the increase in Dpp signalling, as assayed by pMad immunostaining, is concomitant with mitotic entry. Conversely, activation of Dpp signalling in G2 arrested cells does not correlate with elevated levels of pMad. How Dpp signalling is regulated in tracheoblasts is currently unclear. Future studies on how Wnt signalling and Chk1 expression impact Dpp signalling can illuminate how these two signalling pathways act in concert to orchestrate tracheoblast growth and proliferation during larval life.

## ACKNOWLEDGEMENTS

We thank Shigeo Hayashi for fly strains and the Central Imaging and Flow Cytometry Facility (CIFF) and Fly Facility at BLiSc for their support. We thank Benny Shilo, members of the Regulation of Cell Fate Theme at inStem and Narmada Khare for comments on the manuscript. Support: Ramalingaswami Fellowship (Department of Biotechnology, Government of India, AG) and institutional funds from inStem (AK, RSK, PV).

## MATERIALS AND METHODS

### Fly Strains and Handling

The following strains were obtained from repositories: *TubGAL80*^*ts*^; *TM2/TM6b,Tb, UAS-FUCCI, UAS-Fz2*^*RNAii*^, *UAS-Wnt5*^*RNAi*^, *UAS-Wnt6*^*RNAi*^, *UAS-Wnt10*^*RNAi*^, *UAS-CyclinB*^*RNAi*^, *Nkd-LacZ, Chk1-Lacz, Fz3-GFP, UAS-ArmS10, UAS-Tkv*^*QD*^, *Wnt6 KO, Wnt5*^*400*^, *Wg*^*ts*^ (Bloomington Drosophila Stock Center), *UAS-Chk1*^*RNAi*^, *UAS-TCF*^*RNAi*^, *UAS-Wg*^*RNAi*^, *UAS-Dsh*^*RNAi*^, *UAS-Arm*^*RNAi*^, *UAS-Cdc2*^*RNAi*^, *UAS-Tkv*^*RNAi*^, *UAS-Stg*^*RNAi*^ (Vienna Drosophila Resource Center), *UAS-Chk1* (In-house fly facility). The following strains were received as gifts: *Btl-Gal4*. Strains were raised on a diet of cornmeal-agar and maintained at 25°C except GAL80^ts^ strains that were maintained at 18°C. For experiments involving GAL80^ts^ strains, the animals were moved to 29°C at indicated stages for indicated time periods.

### Larval Staging

Larval staging was performed as previously described(Guha and Kornberg, 2005) based on the morphology of the anterior spiracles. L2 larvae were collected and examined to identify animals that had undergone the L2-L3 molt in 8 hour intervals (0-8 h L3). 0-8 h L3 cohorts collected in this method were staged for subsequent time points.

### Immunohistochemistry and Imaging

Animals were dissected in PBS and fixed for 30 minutes with 4% (wt/vol) Paraformaldehyde in PBS. The following antisera were used for Immunohistochemical analysis: Chicken anti-GFP (Aves, 1:500), Rabbit anti-phospho Chk1 (CST, 1:200), Rabbit anti-phospho Smad (CST, 1:150) Rabbit anti-pH3 (Millipore, 1:500), and Alexa 488/568/647-conjugated Donkey anti-Chicken/Rabbit/Mouse secondary antibodies (Invitrogen, 1:200). Tyramide signal amplification was used as per manufacturer recommendations for p-Chk1 detection. The following reagents were used as part of this protocol: Tyramide amplification buffer and Tyramide reagent (Invitrogen), Vectastain A and B and Biotinylated donkey anti Rabbit IgG (1:200, Vector Labs). Tracheal preparations were flat-mounted in ProLong Diamond Antifade Mountant with DAPI (Molecular Probes) and imaged on Zeiss LSM-780 laser-scanning confocal microscopes. Images were processed using Image J. For quantification of cell number, fixed specimens were mounted in ProLong Diamond Antifade Mountant with DAPI and the number of nuclei were counted on an Olympus BX 53 microscope. The DT of the second thoracic metamere was identified morphologically based on the cuticular banding pattern at anterior and posterior junctions.

### Single molecule FISH (smFISH)

Probe sets for *Wg, Wnt5, Wnt6* and *Wnt10* were designed using Stellaris RNA FISH Probe Designer (Biosearch Technologies, Inc., Petaluma, CA) available online at www.biosearchtech.com/stellarisdesigner. smFISH was performed as per manufacturers instructions. Briefly, the tissues were fixed for 45 minutes in 4% (wt/vol) paraformaldehyde at room temperature and permeabilized overnight in 70% ethanol. The hybridization was performed overnight at 37°C. The samples were then washed and imaged on a Zeiss LSM-780 laser-scanning confocal microscope. The probes for *Wg* and *Wnt6* were validated by detecting the pattern of the stripe in WL3 wing disc. *Wnt5* probes were validated by staining the trachea from a *Wnt5* null mutant where we did not observe any signal.

### RNA isolation and quantitative PCR

RNA extraction and qPCR were performed as described in (Kizhedathu et al., 2018). Primer sequences for *Chk1, Fz3, Wg, Wnt5, Wnt6, Wnt10* and *GAPDH* (internal control) are provided below. Relative mRNA levels were quantified using the formula RE= 2^-ΔΔCt^ method.

The following primer sets were used:

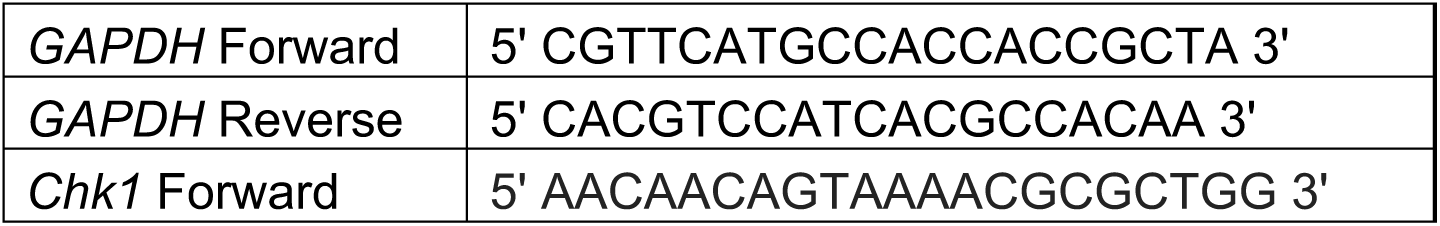
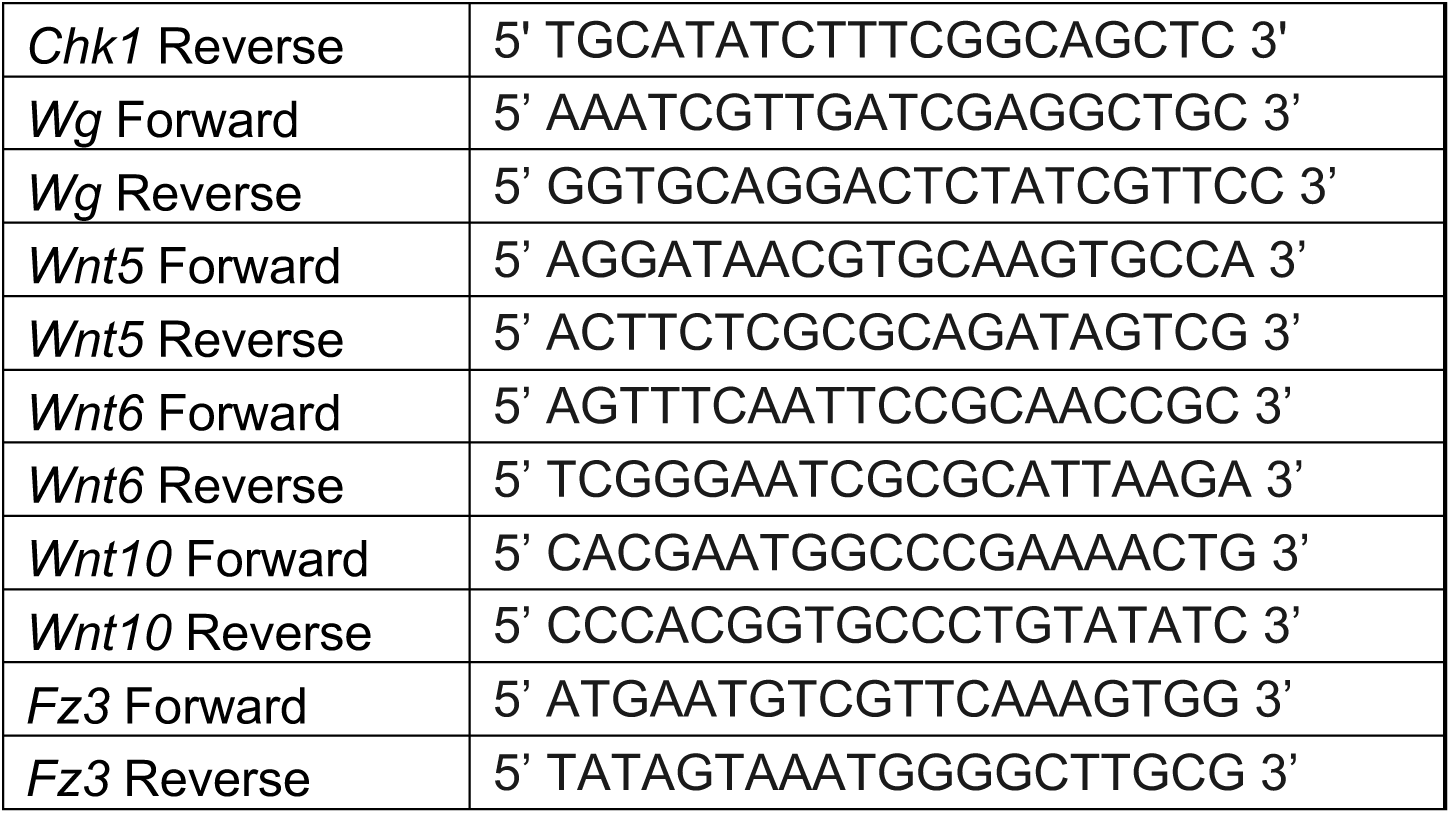

