## SupplementaryFigures for "Multiple Wnts act synergistically to induce Chk1/Grapes expression and mediate G2 arrest in Drosophila tracheoblasts"

**Figure 1 - Figure Supplement 1: Loss of Chk1 or Wnt signalling leads to continuous proliferation of tracheoblasts through L3. (A)** Impact of knockdown of *Chk1* and *TCF* on cell numbers in Tr2 DT. Graph shows cells numbers in wild type (*Btl-Gal4*), *Btl-Chk1*$^{RNAi}$*,* and *Btl-TCF*$^{RNAi}$ at L2, 0-8 h L3, 16-24 h L3 and 32-40 h L3. (L2 and 16-24 h L3 data same as Figure 1B, mean values±standard deviation, n>7 tracheae) Student's paired t-test: *p<0.05

**Figure 5 - Figure Supplement 1: Loss of Stg, Cdc2 or Cyclin B perturbs hypertrophic growth of Tr2 DT.(A)** Impact of reduction of Chk1 or different drivers of G2-M on the size of Tr2 DT at 32-40 h L3. Scatter plot shows length and width of Tr2 DT from Wild type, *Btl-Chk1*$^{RNAi}$*, Btl-Stg*$^{RNAi}$*, Btl-Cdc2*$^{RNAi}$ and *Btl-CyclinB*$^{RNAi}$ at 32- 40 h L3. Values shown in red are from data previously published in (Kizhedathu et al., 2018). (n≥6 tracheae)

### Figure 1 - Figure Supplement 1

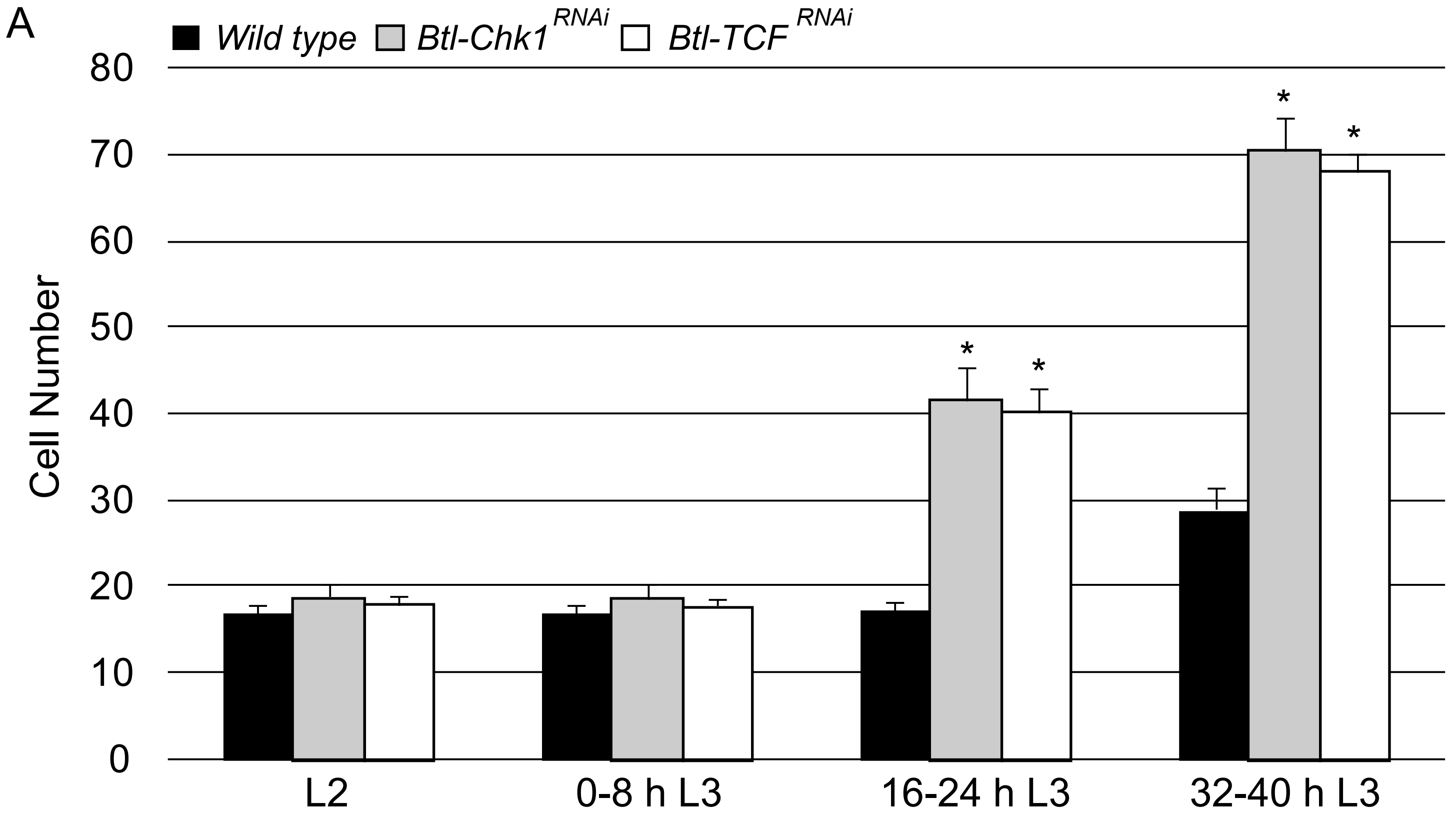

### Figure 5 - Figure Supplement 1

A

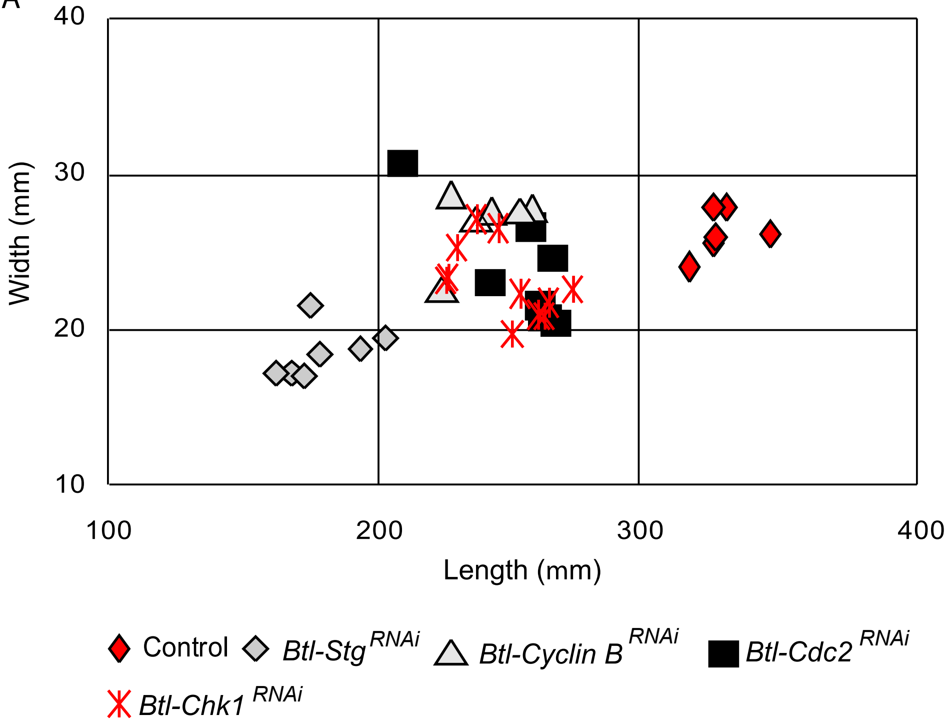
